# Rac1/ROCK-driven membrane dynamics promote Natural Killer cell cytotoxicity via necroptosis

**DOI:** 10.1101/867127

**Authors:** Yanting Zhu, Jun Xie, Jue Shi

**Affiliations:** Center for Quantitative Systems Biology, Department of Physics and Department of Biology, Hong Kong Baptist University, Hong Kong, China

## Abstract

Natural Killer (NK) cells play an important role in cancer immunosurveillance and therapy. However, the target selectivity of NK cell activity is still poorly understood. Here we used live-cell reporters to unravel differential epithelial cancer target killing by primary human NK cells. We found highly variable fraction of killing by distinct NK cell cytotoxic modes that was not determined by NK ligand expression. Rather, epithelial plasma membrane dynamics driven by ROCK-mediated blebs and/or Rac1-mediated lamellipodia promoted necrotic mode in preference to apoptotic mode of killing. Inhibition of granzyme and key necroptosis regulators RIP1, RIP3 and MLKL significantly attenuated necrotic killing, revealing a novel NK cytotoxic pathway by granzyme-induced necroptosis that conferred target selectivity. Our results not only elucidate a new NK cell effector mechanism but also suggest tissue microenvironment and oncogenic signaling pathways that promote membrane dynamics, e.g., Rac1 and Rho/ROCK, could be exploited to enhance proinflammatory NK cell killing.

## Introduction

Natural Killer (NK) cells detect and kill virus-infected cells and cancer cells, and are considered a promising immunotherapeutic candidate [1, 2]. Harnessing and stimulating the direct cytotoxic activity of NK cells has attracted growing interest in the recent development of cancer immunotherapy, as diverse tumor cell types showed susceptibility to NK cell killing and targeted therapy and immune checkpoint therapy may also act in part via triggering NK cell response [3–5]. Tumors are known to harbor large genetic and phenotypic heterogeneity that likely impact their differential interactions with NK cells. However, the mechanistic basis underlying the dynamic control over NK cells’ selective killing of distinct target cell types, e.g., normal vs. cancer as well as different types of cancer, such as those with hot vs. cold inflammatory microenvironment, is poorly understood.

Variable sensitivity of cancer cells to killing by NK cells can be broken into two conceptually distinct processes, detection of abnormal cancer targets, and selection of killing mechanism from the NK cell arsenal, which becomes important if the cancer target is resistant to one or more killing pathways. Determinants of overall effectiveness of NK cells towards particular cancer targets, in terms of both the extent of cytotoxic response as well as cytotoxic mechanism, is still poorly understood. So far the field has very much focused on molecular determinants, while ignoring possible cell mechanical determinants.

Comparatively, target detection has been much more studied than the choice of killing mode. NK cells are known to detect abnormal cells by integrating signals from various inhibitory and activating receptors on their surfaces [6, 7]. The major inhibitory receptors on NK cells are killer cell immunoglobulin-like receptors (KIRs), which interact with major histocompatibility complex I (MHCI) molecules of the target cells, leading to NK cell “self-tolerance” but activation when encountering “foreign” or transformed targets, such as tumor cells with low MHCI expression. Among the many different NK cell activating receptors, NKG2D has been particularly implicated in NK cell-mediated immunity against tumors (8).

Ligands for NKG2D, such as MICA (MHC class I-related gene A), MICB (MHC class I-related gene B) and ULBP (UL16-binding protein), are expressed in most human tumors [9], and inhibiting NKG2D function attenuated NK cell cytotoxicity against tumor targets [10].

The mechanisms underlying choice of killing mode could depend on sensitivity of the targets towards different modes and/or triggering of alternative cytotoxic pathways in the NK cells. NK cells are thought to kill targets by mechanisms including lytic granule-mediated apoptosis, death ligand-mediated apoptosis, pyroptosis and necrosis [11–15]. Lytic granule-mediated apoptosis is considered the principal cytotoxic pathway of NK cells. Upon formation of NK-target cell immunological synapses, lytic granules release perforin and granzymes into the synapse. Granzymes then enter the target cell through the perforin channels, where they activate the apoptosis pathway via proteolytic activation of caspases [16]. In addition, NK cells express a variety of death ligands, such as Fas ligand (FasL) and TNF-related apoptosis inducing ligand (TRAIL), which engage cognate death receptors expressed on the target cell surface, triggering death by receptor-mediated extrinsic apoptosis [17, 18]. Two recent studies identified pyroptosis as a new NK cell cytotoxic mechanism, in particular activating proinflammatory cell death in tumors that express gasdermin B or gasdermin E [13, 14]. Another highly proinflammatory NK cell killing mode is necrosis, which has been reported in a few studies and its mechanistic basis is still largely unknown [15, 19]. Here, we identified necroptosis as an NK cell cytotoxic effector mechanism for the first time. It may be a distinct cytotoxic mode, or equivalently underlying the necrotic killing that was previously reported. In light of the strong antitumor activity of NK cell-induced pyroptosis due to its proinflammatory consequences, the elucidation of necroptosis as a new proinflammatory NK cell killing mode is exciting, as this killing mode is not restricted by gasdermin expression and may be exploited for more diverse tumor subtypes.

To unravel the target selectivity of NK cell cytotoxicity, in this study we characterized the rate-limiting kinetics and phenotypic heterogeneity of NK cell cytotoxic activity against a panel of normal and epithelial cancer targets using a live-cell FRET (Förster Resonance Energy Transfer) reporter and quantitative single cell imaging. Sensitivity of the epithelial target cells to overall NK cell killing was mainly conferred by the inhibitory KIRs-MHCI interaction, as expected. However, the choice of cytotoxic mechanism, e.g., necrosis vs. apoptosis, was determined by differential plasma membrane dynamics of the epithelial cell targets. In particular, a highly dynamic target membrane driven by Rac1 and/or Rho/ROCK activities, which are responsible for membrane protrusion and contractility, promoted granzyme-induced actin depolymerization followed by target membrane leakage and formation of large blebs, ultimately leading to target cell death by the highly proinflammatory necroptosis. The identification of granzyme-induced necroptosis as a new cytotoxic effector mechanism revealed an alternative signaling cascade to understand cytotoxic lymphocyte activity as well as necroptosis regulation in other important developmental and disease processes.

## Results

### Target cell-type dependence of NK cell cytotoxicity

To investigate mechanisms underlying the differential activity of NK cells on distinct targets, we chose a panel of human epithelial cell lines that exhibited variable sensitivity to primary NK cell killing, including one normal cell line, LO2 (immortalized normal hepatic cell line) and four cancer cell lines, i.e., HeLa (cervix cancer), SMMC-7721 (liver cancer), MCF7 (breast cancer), and U-2 OS (bone cancer). To visualize dynamic response to NK cells, we engineered each epithelial cell line with a fluorescent FRET reporter of granzyme-B activity (GzmB-FRET) as well as a mitochondria reporter of apoptosis (IMS-RP), as reported previously [20–22]. All NK-target cell co-culture assays were conducted with human primary NK cells purified from healthy blood donors and pre-activated by IL-2 for three days, at an NK-to-target cell ratio of 3:1. Target cell death was scored morphologically by blebbing followed by cell lysis, and the time from NK cell addition to morphological target cell death was plotted as cumulative survival curves. As shown in Figure 1A, the normal cell line, LO2, was the most resistant to NK cell killing, consistent with NK cell’s function in eliminating abnormal and cancerous cells. Among the cancer cell lines, SMMC-7721 and MCF7 were the most sensitive, with MCF7 exhibiting the fastest kinetics to cell death.

**Figure 1.**
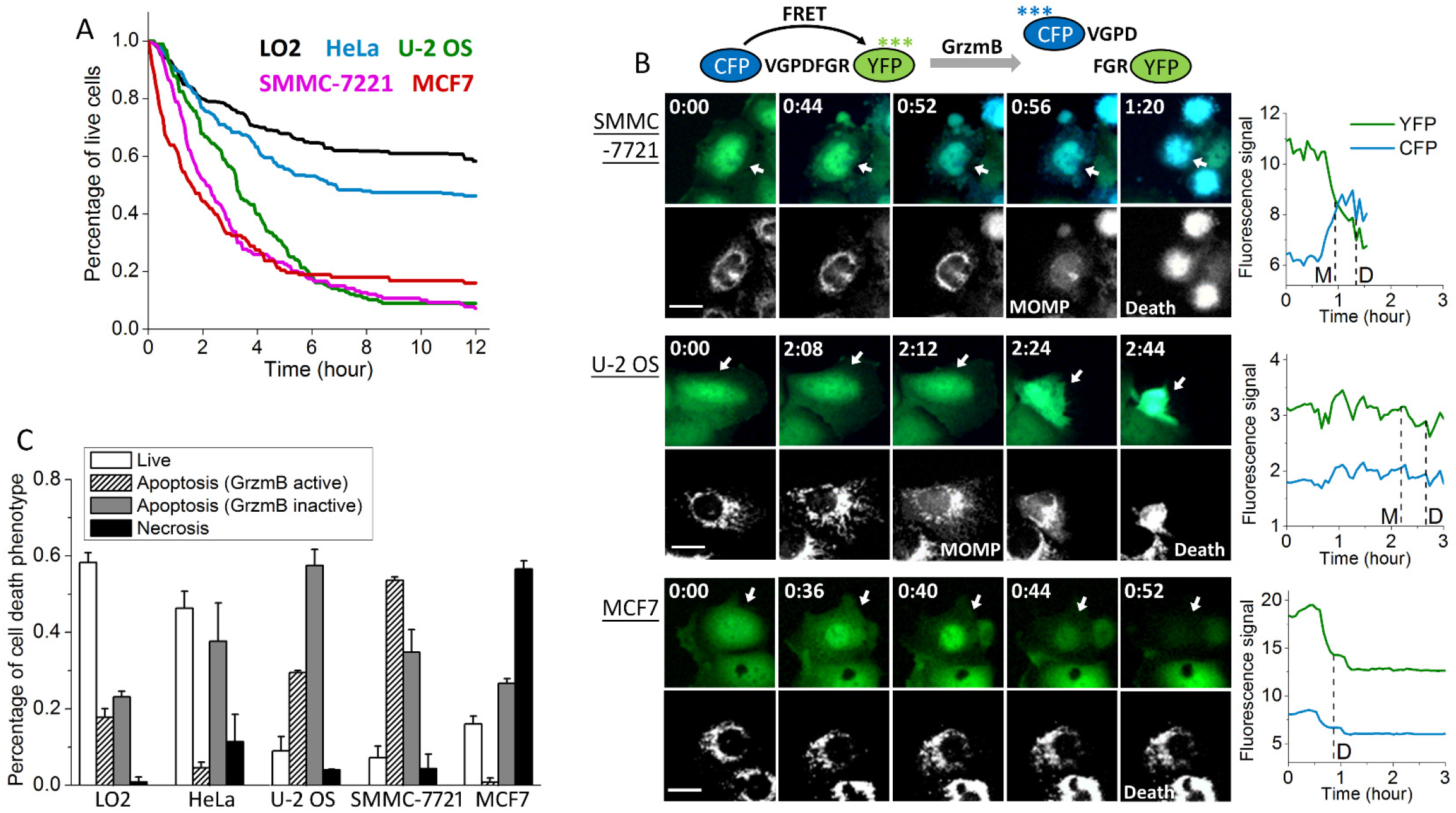
Characteristics of the cytotoxic dynamics of primary NK cell against five different epithelial target cell lines. (A) Cumulative survival curves of target cells, including LO2 (denoted in black), HeLa (green), U-2 OS (blue), SMMC-7721 (magenta) and MCF7 (red), in co-culture with primary human NK cells. (B) Left panel: Fluorescent images of the granzyme-B FRET (GzmB-FRET) reporter and the corresponding mitochondria reporter, IMS-RP, from SMMC-7721, U-2 OS and MCF7, respectively. The GzmB-FRET images are overlay of the CFP (denoted by blue) and YFP (green) channels. Time (unit: hour:minute) is indicated at the top left corner of each GzmB-FRET image. Right panels: Single cell trajectories of CFP and YFP signals quantified from the time-lapse movies. The time of MOMP (scored by the IMS-RP signal change) and the time of death (scored morphologically by cell blebbing and lysis) are indicated by the vertical dotted line. The white scale bar is 20 μm. (C) Distributions of the live and dead target cells killed by the three distinct cytotoxic modes mediated by granzyme B (GrzmB active), death ligand (GrzmB inactive) and necrosis after 12 hours of co-culture with primary NK cells. Data plotted in (A) and (C) were averaged from 2 independent imaging experiments and the number of cells analyzed ranges from 40 to 223, varied between experiments and target cell lines. The error bars are standard deviations.

In addition to variable sensitivity to overall NK cell killing, we also observed striking difference in the cytotoxic modes used to kill the different target cell lines. The majority of cell death events (about 60%) seen in SMMC-7721 were preceded by a loss of the granzyme-B FRET (i.e., an increase of CFP (donor) signal and decrease of YFP (acceptor) signal) followed by mitochondrial outer membrane permeabilization (MOMP), suggesting that NK cells killed SMMC-7721 mainly by the lytic granule and granzyme-B mediated cytotoxic pathway and intrinsic apoptosis (Fig. 1B). In contrast, NK cell cytotoxicity towards U-2 OS cells mostly did not associate with a change in the granzyme-B FRET signal. We have previously characterized this mode of target cell death to be mediated by the death ligand, e.g., FasL, and subsequent extrinsic apoptosis [21]. Interestingly, we found necrosis, a largely unexplored NK cell cytotoxic mechanism, was the primary cytotoxic mode that triggered cell death in MCF7 cells. As shown in the bottom panel of Figure 1B, MCF7 cell death was not preceded by MOMP, indicating it was not classic apoptosis. Moreover, signal from the granzyme-B FRET reporter was abruptly lost upon extensive MCF7 cell blebbing, pointing to large scale leakage of the intracellular content likely due to membrane ruptures. Such dynamic features were consistent with necrotic cell death.

Figure 1C summarized the percentage of live target cells and the cell death population via the three distinct cytotoxic modes of primary NK cell exhibited by the five epithelial target cell lines after 12 hours of NK-target cell co-culture. The two most sensitive target cell lines, SMMC-7721 and MCF7, which showed rapid kinetics of cell death induction, were killed mainly (around 60%) through granzyme-B activity or necrosis, while cytotoxicity mediated by death ligand was more dominant in the less sensitive target cell types, U-2 OS, HeLa and LO2. All target cell lines showed substantial cell death triggered by death ligands of NK cells, ranging from 23% to 57% of the total target cell population, suggesting the death receptor pathway is widespread. In contrast, the extent of cell death activated by the granzyme-B and necrosis pathways varied more significantly, ranging from 1% to 54% for the lytic granule mode and 1% to 57% for the necrotic mode. Such large variability in the sensitivity to NK cell killing via granzyme-B and necrosis indicated that activation of these two cytotoxic modes may depend on epithelial features that are more cell type specific.

### Extent of cytotoxicity, but not cytotoxic mode, correlated with MHCI expression

To investigate the potential molecular determinants underlying the observed variable sensitivity of epithelial cell lines to overall NK cell killing as well as to the distinct NK cell cytotoxic modes, we first profiled the expressions of NK cell-interacting surface molecules known to be involved in cancer-associated NK cell cytotoxicity for the selected target cell panel (Fig. 2A). These molecules include the human MHCI molecule (HLA-A,B,C), NKG2D ligands (MICA, MICB and ULBP-2,5,6) and DNAM-1 ligands (PVR/CD155 and Nectin-2/CD112) (21). We also measured the expression of the death receptor Fas and an integrin key for NK cell adhesion, ICAM-1 (CD54), in our analysis by flow cytometry (Fig. 2B). The quantified FACS results showed highly variable expressions of all surface molecules that we profiled between the five target cell lines (Fig. 2C). The sensitivity of target cell lines to overall NK cell killing correlated relatively well with the expression level of the inhibitory MHCI molecules, and to a much lesser extent with MICB, but did not correlate with the expression of the other NKG2D activating ligands, MICA and ULBPs, or the activating ligands for DNAM-1. Surprisingly, the expression level of Fas did not correlate with the differential sensitivity to the death ligand-mediated NK cell cytotoxicity. Our data thus indicated that the inhibitory strength of KIRs-MHCI may exert the primary control over target cell recognition and overall NK cell cytotoxicity against a particular epithelial target, while control by the activating ligands is less universal and more context-dependent. However, our data did not show any significant correlative feature that was specific to the three distinct NK cell cytotoxic modes. The only differential ligand expression that may potentially render specificity to necrotic killing is MICB, as the MICB expression level in MCF7, the cell line that was particularly prone to necrotic death, was 5-10 folds higher than that in the other four cell lines. However, RNAi knockdown of MICB in MCF7 cells did not attenuate necrotic death induced by NK cells (data not shown), indicating MICB expression was not the molecular determinant for the necrotic killing.

**Figure 2.**
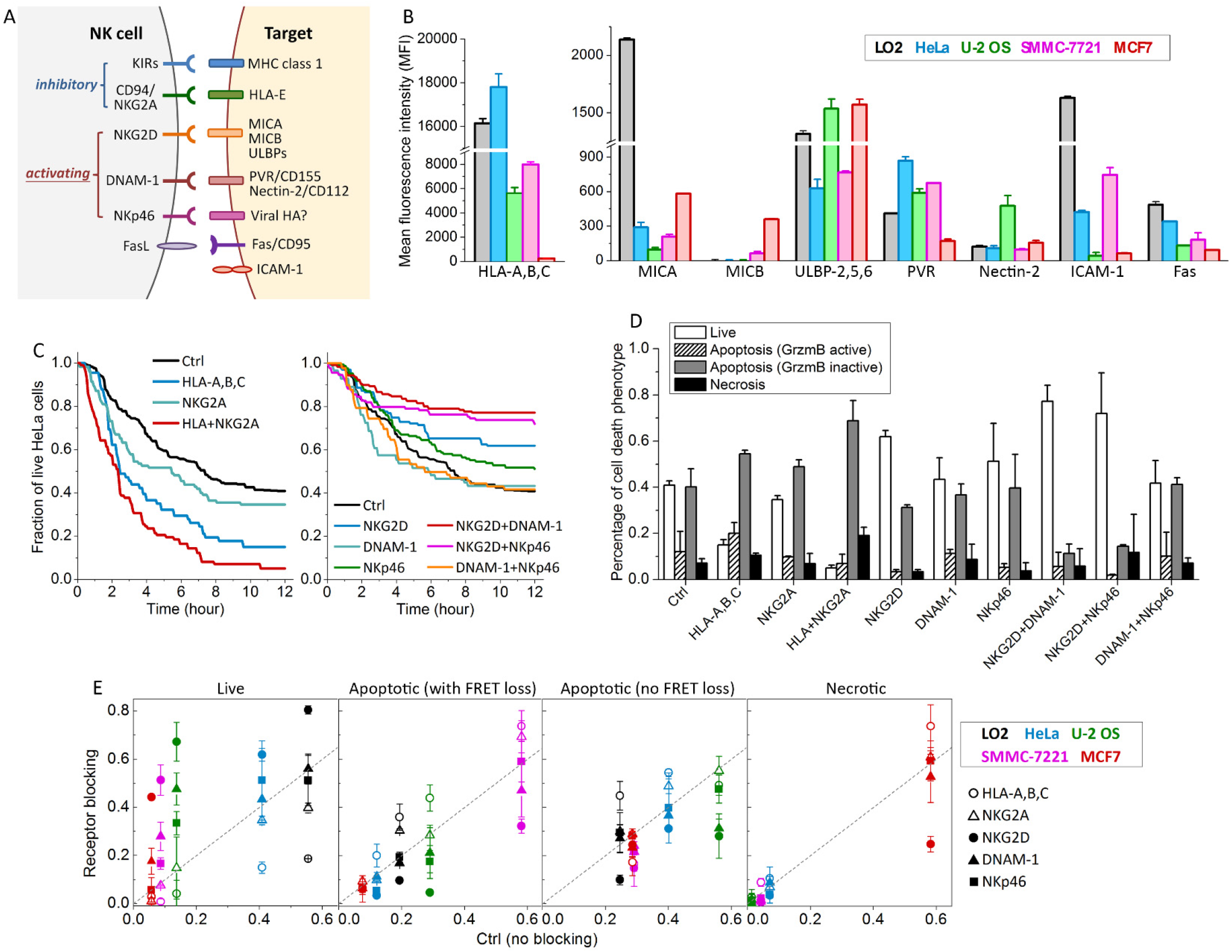
Differential activation of the NK cell inhibitory and activating receptor signaling machinery upon interaction with the five epithelial target cell lines. (A) Diagram of the major NK cell receptors and their cognate ligands on the target cell. (B) Comparison of the surface protein expression in the five target cell lines based on the average florescent signal from the flow cytometry analysis. Cell lines were color coded as indicated. Data were averaged from 2 independent flow cytometry analyses (>1×10^4^ cells in each analysis). The error bars are standard deviations. (C) Cumulative survival curves of HeLa cells in co-culture with primary NK cells in the presence of single or double blockage of the inhibitory receptor (left panel) or activating receptor (right panel). (D) Distributions of the live and dead HeLa cells killed by the three distinct cytotoxic modes after 12 hours of co-culture with NK cells under the indicated treatment conditions. (E) Percentage of live cells and cells that died via the three cytotoxic modes under the different receptor blocking conditions in comparison with those under the control condition. Data from the five target cell lines were color coded as indicated. The different treatment conditions were denoted with the indicated symbols. Data plotted were averaged from 2 or 3 independent imaging experiments and the error bars are standard deviations.

As we did not observe correlation between the expressions of well-known NK cell-interacting surface ligands with the variable sensitivity to different NK cell cytotoxic modes, we went on to investigate the involvement of key inhibitory and activating receptors on NK cell surface, as the target specificity could be conferred by surface ligands beyond the profiling panel that we selected. The NK cell receptors that we examined included two types of inhibitory receptors, KIRs and NKG2A (CD94), and three activating receptors important for cancer target recognition, including NKG2D (CD314), DNAM-1 (CD226) and NKp46 (CD335) (11, 21). Specifically, we used neutralizing antibodies to block individual receptors or combination of the receptors and then compared the cytotoxic response via the three cytotoxic modes with those under the control condition (i.e., no receptor blocking). And we used a broad-spectrum neutralizing antibody against HLA-A, -B and -C to block the interaction of all KIRs and human MHCI molecules as a whole, instead of examining the individual KIR, to simplify the analysis.

Figure 2C and 2D showed the receptor neutralizing results for HeLa cells in co-culture with primary NK cells. As expected, blocking the inhibitory NK-HeLa cell interaction via MHCI molecules or NKG2A accelerated and enhanced HeLa cell killing, with MHCI neutralization showing a stronger effect (Fig. 2C). Simultaneous blockage of MHCI molecules and NKG2A further increased NK cell cytotoxicity towards HeLa, resulting in a degree of cell death similar to that observed in the sensitive target cell lines, such as SMMC-7721. In contrast, neutralizing the activating receptors exerted less prominent effect in attenuating the cell death response of HeLa, possibly because HeLa cells under the control condition were already relatively resistant to primary NK cell killing. Inhibition of NKG2D activity exhibited a stronger effect in attenuating cell death than neutralizing DNAM-1 or NKp46 (Fig. 2C). Double blocking of NKG2D plus DNAM-1 or NKp46 further rescued HeLa cell death, confirming that the NK cell cytotoxicity is regulated by collective, rather than individual, signaling receptors.

We next investigated how blocking individual receptors altered induction of the three cytotoxic modes, expecting individual receptors to selectively regulate particular modes. However, this was not the result. The fraction of NK cell killing mediated by granzyme-B, death ligand and necrosis all increased in parallel upon blockage of the inhibitory KIR-MHCI interaction, and decreased in parallel upon neutralizing the activating receptor NKG2D, either alone or in combination with DNAM-1 or NKp46 (except for the necrotic population under NKG2D+ NKp46 inhibition). This parallel activation or inhibition of all three pathways can be seen by the similar relative heights of the three non-white bars in Figure 2D. Therefore, in HeLa, receptor modulation appears to tune the overall cytotoxic activity of NK cells and/or receptivity of targets, but not the specific death pathway.

Figure 2E summarized the receptor neutralization results for all three pathways across the five epithelial target cell lines. Here, we plotted the ratio of perturbation versus control. Intuitively, data points along the diagonal indicated no change relative to the control condition, and the further away the data points were from the diagonal, the larger the effect of the respective receptor inhibition in diverting the cell death response into, or away from, one of the three cytotoxic modes. Similar to HeLa cells, neutralization of the inhibitory KIRs-MHCI interaction exerted strong effect in enhancing cell death response of the other four target cell lines, and loss of NKG2D activity exerted the strongest effect in attenuating target cell death. Double inhibition of NKG2D and DNAM-1 nearly abrogated cell death of U-2 OS, SMMC-7721 and MCF7, the three cell lines that were most sensitive to NK cell killing. However, as of HeLa cells, the three cytotoxic modes were altered approximately in parallel, again showing that receptor modulation tuned overall activity of NK cells, but not the activity of one specific death pathway. We also noted that except for MCF7, no other cell line showed significant cell death via necrosis under all receptor perturbation conditions, suggesting that MCF7 cells may have unique cellular features that promote the induction of necrotic killing, which we further investigated below.

### Membrane dynamics modulate target sensitivity to NK cell cytotoxic modes

Since receptor expression level did not predict which death pathway is preferred, and receptor inhibition did not modulate the fraction of death caused by particular pathway, we sought other phenotypic properties that might predict and modulate individual NK cell cytotoxic pathways. We focused on MCF7 and SMMC-7721 because they had the most distinctive preferences, i.e., MCF7 cells were uniquely sensitive to necrosis, while SMMC-7721 cells were sensitive to granzyme-B-mediated intrinsic apoptosis. A distinguishing feature of these cell lines that we observed in our movies was differences in plasma membrane dynamics. MCF7 cells exhibited extensive dynamic protrusions driven by membrane blebs and lamellipodia (Fig. 3A). In contrast, SMMC-7721 cells exhibited only moderate membrane dynamics and the membrane of U-2 OS cells was largely quiet (Fig. 3A). This observation led us to examine whether differential membrane dynamics of the epithelial cell targets contributed to the activation of alternative NK cell cytotoxic modes.

**Figure 3.**
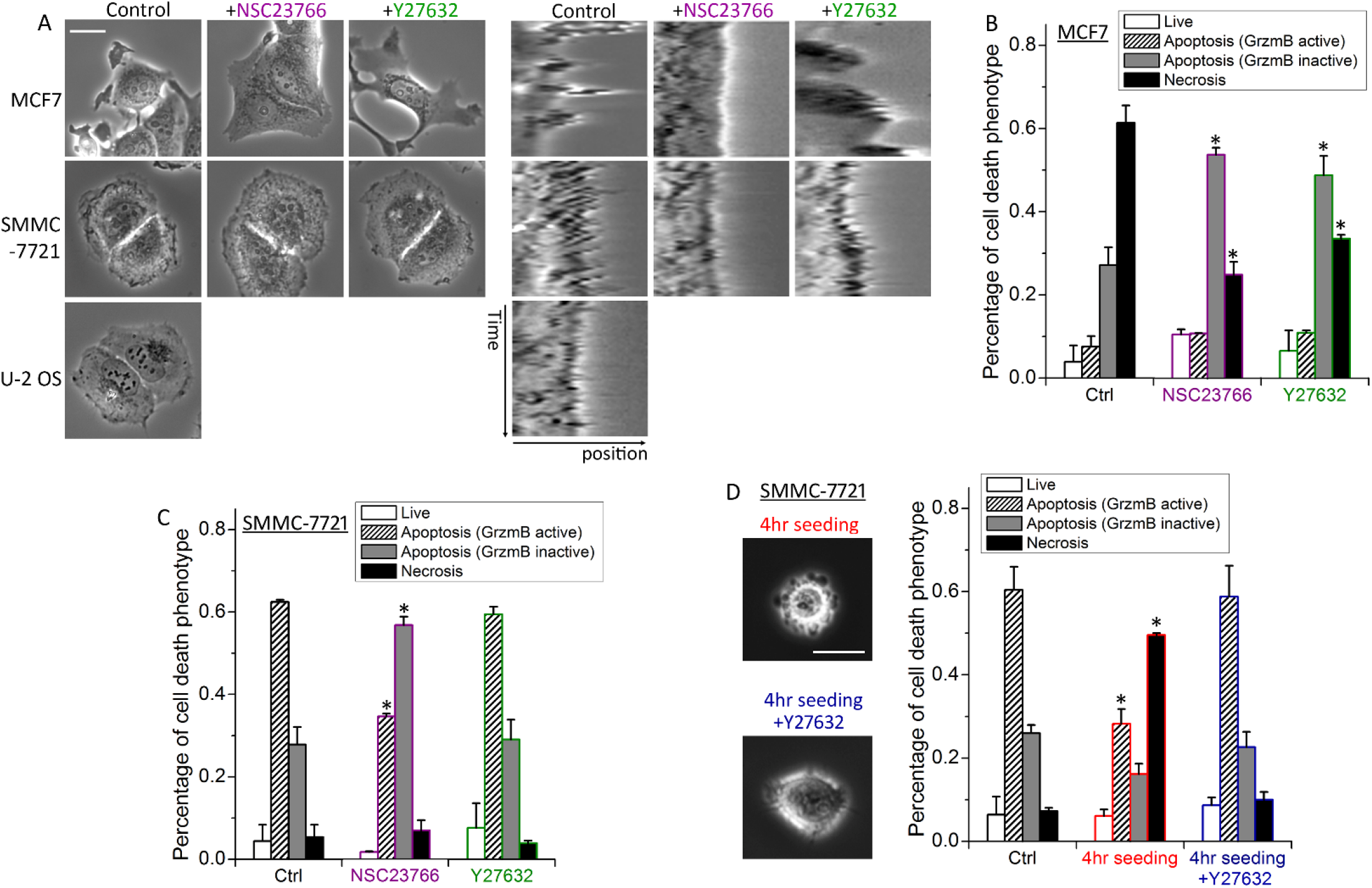
Membrane dynamics of MCF7 and SMMC-7721 cells modulate NK cell cytotoxic modes. (A) Left panel: Phase-contrast images of MCF7, SMMC-7721 and U-2 OS cells under control or treatment with NSC23766 or Y27632. Right-panel: Kymograph of a cell edge of MCF7, SMMC-7721 and U-2 OS cells under the indicated treatment conditions. The white scale bar is 20 μm. (B, C) Distributions of the cytotoxic response phenotypes of target cells under control or the indicated inhibitor treatment for (B) MCF7 cells or (C) SMMC-7721 cells in co-culture with primary NK cells for 12 hours. (D) Phase-contrast images and the response phenotype distribution of SMMC-7721 cells that adhered to the cell culture plate for 4 hours (with or without Y27632) and in co-cultured with primary NK cells for 12 hours in comparison with control cells that adhered for 24 hours. Data plotted in (A) to (D) were averaged from 2 to 4 independent imaging experiments and the number of cells analyzed for each condition/cell line/experiment ranges from 44 to 149. The error bars are standard deviations. P value was obtained by student’s t test comparing the treatment condition with control. * P < 0.001.

To attenuate the formation of dynamic membrane protrusions, we used either NSC23766, an inhibitor of the small GTPase Rac1, to inhibit actin polymerization-driven lamellipodia, or Y27632, an inhibitor of the Rho Kinase (ROCK), to inhibit actomyosin contractility that promotes membrane blebs [24–26]. NSC23766 significantly reduced membrane protrusions in both MCF7 and SMMC-7721 cells, as evidenced by the kymograph of cell edge, which showed a relatively smooth edge under NSC23766 as compared to a jagged edge in the control cells (Fig. 3A). The effect of Y27632, however, diverged in MCF7 and SMMC-7721. MCF7 cells treated with Y27632 showed reduced membrane blebs but longer lamellipodia, a phenotype previously characterized as due to compensatory effects of membrane blebs and lamellipodia [24]. SMMC-7721 treated with Y27632 still showed evident membrane ruffling, indicating bleb did not contribute significantly to drive the membrane dynamics of SMMC-7721. Attenuation of membrane dynamics by both NSC23766 and Y27632 changed the primary cell death mode of MCF7 cells from necrosis to death ligand-mediated apoptosis, with NSC23766 exhibiting a slightly stronger effect (Fig. 3B). NSC23766 treatment of SMMC-7721 also resulted in a switch of NK cell killing from granzyme-B-mediated apoptosis to death ligand-mediated apoptosis, while Y27632 did not show significant effect on altering the NK cell cytotoxic mode (Fig. 3C). Our data thus showed inhibiting plasma membrane dynamics of epithelial targets promoted NK cell killing via death ligand-mediated apoptosis, while damping the other two cytotoxic pathways.

Next we examined whether increasing membrane dynamics, e.g., in SMMC-7721, would alter NK cell killing from graznyme-B-mediated apoptosis to the necrosis mode, similar to MCF7. We noted that SMMC-7721 cells showed extensive membrane blebs when they started to adhere to the cell culture plate surface (Fig. 3D, left panel). And these blebs disappeared upon complete adherence (Fig. 3A, control). We therefore co-cultured primary NK cells with SMMC-7721 that were trypsinized and seeded onto the culture plate for only 4 hours to investigate the effect of enhanced membrane dynamics, in this case, driven by blebs, on NK cell killing mode. Compared to control SMMC-7721 cells that had adhered to the culture plate for 24 hours, which were mainly (~60%) killed by granzyme-B-mediated apoptosis, the majority (~50%) of the short-adherence cells were killed via the necrotic mode (Fig. 3D, right panel). And addition of the bleb inhibitor, Y27632, abrogated the increase in necrotic killing of the short-adherence cells (Fig. 3D). As U-2 OS cells adhered to the culture plate surface very quickly (in about 3-5 hours) and did not exhibit prolonged blebs, we were unable to use this experimental strategy to study the effect of enhanced membrane dynamics on promoting granzyme-B-mediated apoptosis over the death ligand-mediated mode. Nonetheless, overall our results suggested that the extent of membrane dynamics exerted a key control over the sensitivity of different epithelial targets to the three distinct NK cell cytotoxic modes.

### Pro-necrotic membrane dynamics involve actin depolymerization

To determine how a highly dynamic target membrane promotes pro-necrotic killing, we stained the lytic granules of NK cells with an acidic organelle marker, Lysobrite, and investigated the collective dynamics of NK and target cells upon formation of the immunological synapse. NK cells typically showed polarized morphology with the acidic lytic granules stored in the tail end, as they moved in the co-culture environment. Such localization of lytic granules likely prevents undesirable leakage of lytic granules during the constant transient contacts between NK and target cells. When NK cells recognized an abnormal MCF7 cell target, a sustained conjugation called immunological synapse (IS) was formed and subsequently triggered the reorientation of lytic granule from the tail to the IS and then lytic granule transfer (Fig. 4A, left panel; Movie S1). We found localization and transfer of lytic granule at IS was accompanied by large membrane blebs of MCF7 cell, which, within 4-8 minutes, ruptured the MCF7 cell membrane and caused necrotic death. We also noted the large membrane blebs tended to form at sites previously showing large lamellipodia (Fig. 4A, frame #1 and #2), suggesting possible conversion of lamellipodia to bleb. SMMC-7721, in comparison, exhibited significantly less membrane bleb upon the localization and transfer of lytic granules at the IS that was followed by gradual accumulation of granzyme-B activity as indicated by the loss of GrzmB-FRET signal, and induced apoptotic death in 30-40 minutes (Fig. 4A, right panel; Movie S2). Attenuating MCF7 membrane dynamics by inhibiting Rac1 or ROCK activity abrogated the induction of large membrane bleb at the IS and subsequent necrotic killing (Fig. 4B), while increasing the membrane dynamics of SMMC-7721 by short adherence induced a large bleb at the IS and led to necrotic death, similar to the dynamic phenotype that we observed for MCF7 cells (Fig. 4B, right panel).

**Figure 4.**
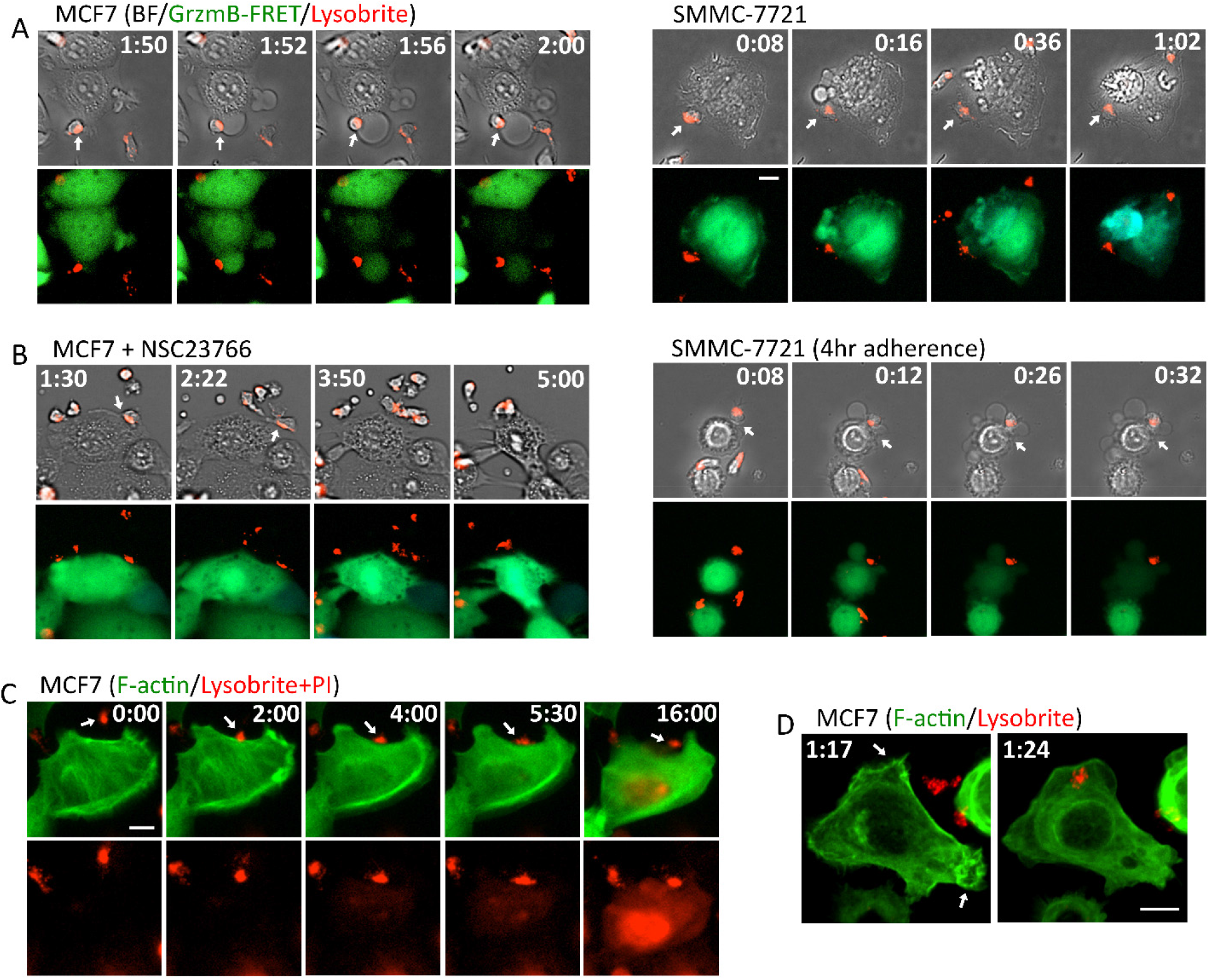
Necrotic killing of NK cells are associated with actin depolymerization and membrane leakage. (A) Representative still images obtained from live-cell imaging of MCF7 (left panel) and SMMC-7721 (right panel) in co-culture with primary NK cells. The upper panel shows the overlay images of bright-field and the red lytic granule marker, lysobrite. The lower panel shows the overlay images of the GrzmB-FRET reporter and lysobrite. Time is indicated in the unit of hour:minute at the top of the images. The specific interacting NK cell is indicated by the white arrows. (B) Still images of MCF7 cells treated with NSC23766 (left panel) and SMMC-7721 cells upon 4-hour adherence (right panel) in co-culture with primary NK cells stained with lysobrite. (C) Dynamics of F-actin (denoted in green) and membrane leakage probed by PI (red) infiltration into the MCF7 cells (detected by the same fluorescent channel as lysobrite) triggered by NK cells. Time is indicated in the unit of minute:second. The imaging frame rate for this time-lapse experiment is 30 seconds per frame. (D) Confocal livecell imaging of F-actin dynamics upon NK-MCF7 cell interaction. The white arrows point to highly dynamic sites of lamellipodia that converted into blebs upon NK cell interaction. Time is indicated in the unit of hour:minute. The white scale bar in (A) to (D) is 10 μm.

As the large membrane bleb was a key feature preceding necrotic killing, we further investigated its mechanistic origin. The observation of possible conversion of lamellipodia to large bleb led us to examine the dynamics of cytoskeleton, in particular F-actin, by imaging a transiently expressed Utrophin-GFP reporter in the MCF7 cells [27]. We also added propidium iodide (PI) in the live-cell imaging assay to determine the onset of membrane leakage in the necrotic killing process. As shown in Figure 4C, lamellipodia immediately stalled and retracted upon IS formation and the actin cytoskeleton largely depolymerized within 2 minutes. Membrane leakage indicated by PI diffusion into the MCF7 cell was generally observed 1-1.5 minutes subsequent to the change in actin dynamics (the PI fluorescent signal was detected in the same channel as lysobrite). Confocal imaging of the F-actin and lysobrite confirmed the change in actin dynamics and cytoskeleton upon lytic granule localization to the IS at higher spatial resolution (Fig. 4D; Movie S3). Distinct from the MCF7 cells, IS formation in SMMC-7721 led to the alignment of F-actin into long fibers, likely due to increased contractility known in apoptotic cells (Movie S4). Together our data suggested that the lytic granules of NK cells induce actin deploymerization and loss of lamellipodia that initiate bleb formation, especially at the highly dynamic membrane sites. Subsequent membrane leakage after cytoskeleton destruction likely also contributed to enhance the bleb by increasing osmotic pressure, eventually leading to necrotic membrane rupture.

### Necrotic NK cell cytotoxicity is granzyme-induced necroptosis

NK cell-induced necrosis is poorly characterized at the molecular level. As MCF7 cells were killed mostly by this mechanism, it provided a model for probing the mechanistic origin of necrotic killing. To test the involvement of lytic granules, we used Concanamycin A (CMA), an inhibitor of vacuolar type ATPase (V-ATPase) that increased the pH of lytic granules, to disrupt lytic granules in NK cells and inhibit their activities. As shown in Figure 5A, treatment of 10 nM CMA reduced the necrotic killing of MCF7 cells from 53% to 4% after 6 hours of co-culture with NK cells and also abrogated the lytic granule-mediated apoptosis, which confirmed the key role of lytic granules in activating these two cytotoxic pathways. The extent of death ligand-mediated apoptosis was not affected by CMA treatment, as expected. We also used EGTA to chelate Ca^2+^ flux that is crucial for lytic granule transfer. Again we found both the necrotic death and lytic granule-mediated apoptosis in MCF7 cells were significantly decreased (Fig. 5A). Moreover, the substantial loss of necrotic MCF7 cell death was also observed by treating the NK cells with a pan-granzyme inhibitor, 3,4-Dichloroisocoumarin (DCI), illustrating that it is the granzyme activity from the lytic granules that induces the actin depolymerization and bleb formation upon lytic granule transfer to the target cells (Fig. 5A).

**Figure 5.**
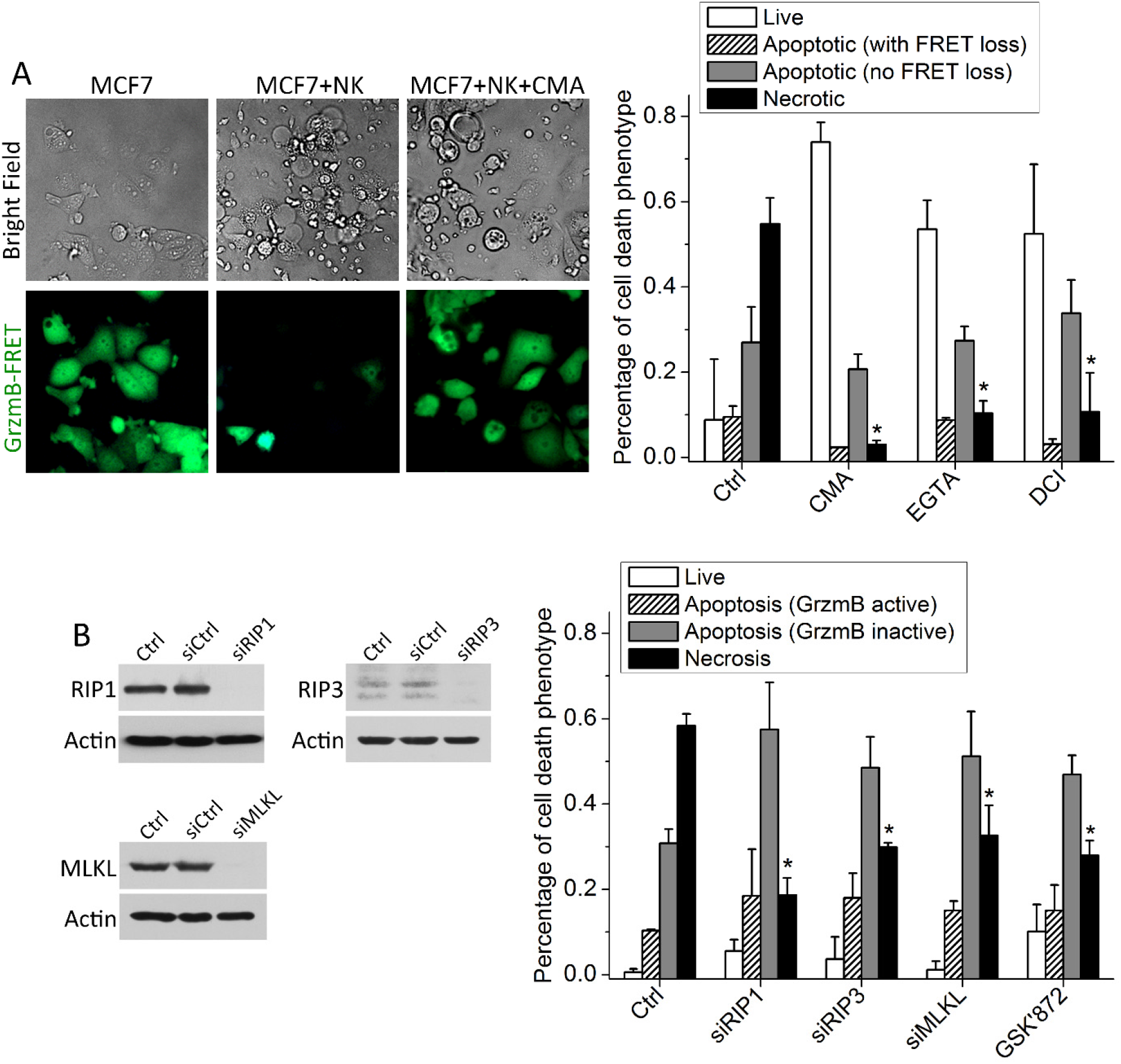
Necrotic NK cell cytotoxicity is triggered by granzyme and mediated by the necroptosis pathway. (A) Left panel: Representative still images of MCF7 cells in co-culture with primary NK cells with or without 10 nM CMA. Right panel: Distribution of the indicated response phenotypes of MCF7 cells after 6-hour co-culture with primary NK cells under the control condition or treatment of 10 nM CMA, 0.9 mM EGTA and 40 μM DCI. (B) Distribution of the cytotoxic response phenotypes of MCF7 cells upon knockdown of RIP1, RIP3 and MLKL, respectively, and under the treatment of 5 μM GSK’872. The knockdown efficiency is demonstrated by the western blots. Data plotted in (A) and (B) were averaged from 2 or 3 independent imaging experiments and the number of cells analyzed for each condition/experiment ranges from 55 to 112. The error bars are standard deviations. P values were obtained by student’s t test comparing the treatment condition with control. * P < 0.0001.

Large membrane blebs dependent on the actin cytoskeleton are characteristic of necroptosis, a type of programmed necrosis that is usually induced by cell surface receptors [28–31]. This pathway has not been previously implicated in the killing by cytotoxic lymphocytes. To test its possible involvement, we knocked down key necroptosis pathway components, including RIP1 (receptor-interacting protein 1), RIP3 and MLKL (mixed lineage kinase domain-like), in MCF7 cells by RNA interference (RNAi). Loss of these three regulators all caused a significant reduction in necrotic killing, and a concomitant increase in apoptotic killing (Fig. 5B). We also treated MCF7 cells with commonly used necroptosis inhibitors. The RIP3 inhibitor, GSK872, decreased necrotic MCF7 cell death to a similar degree as the RIP3 knockdown treatment (Fig. 5B). We were unable to test the inhibitory effects of the RIP1 inhibitor, necrostatin-1, and MLKL inhibitor, necrosulfonamide, as these two inhibitors exhibited toxicity to the primary NK cells, causing loss of polarized morphology and significantly reduced cytotoxic activity. Based on the morphology of death, and sensitivity to inhibition of the necroptosis pathway, we conclude that the primary NK cell-induced death mode in MCF7 targets is granzyme-induced necroptosis.

## Discussion

Our study revealed that epithelial cell targets varied not only in their overall sensitivity to killing by primary NK cells, but also in the mode of killing. Overall sensitivity was determined primarily by inhibitory and activating receptor-ligand interactions, consistent with previous literature. However, these interactions did not specify the killing mode. Rather, membrane dynamics of the epithelial targets appeared to play a central role in cytotoxic mode specification. We also demonstrated necroptosis as an effector mechanism in NK cell killing for the first time. These findings have important implications for immunotherapy of solid tumors, and for understanding regulation of necroptosis in other homeostatic and disease processes.

Our measurements of the different NK cell receptor ligands for tuning overall sensitivity suggest that it should be possible, in principle to predict, and eventually modulate, the sensitivity of different cancers to NK cell therapy. However, the relationship between receptor expression and sensitivity is complex, and measurement of one or two ligands will not be sufficient. For example, we found the normal cell line, LO2, which was the most resistant to NK cell cytotoxicity, expressed the highest level of NKG2D ligand, MICA, and also high level of ULPs when compared with the cancer cell lines (Fig. 3B). Previous clinical studies have also showed that high levels of NKG2D ligands were not associated with the extent of NK cell infiltration into the tumors or better prognosis for patients with breast, lung or ovarian cancers [32], suggesting that expressions of NK cell activating ligands likely vary significantly between cancer types and tumor context, thus hindering their stand-alone prognostic potential.

Our main novel finding concerns variability in NK cell killing mode between different epithelial targets, and the cellular determinants of this choice. Killing mode is important because it determines the molecular pathways by which cancer cells might become resistant under selection, and also because it may influence downstream immunological consequences of killing. As necrotic death releases cell debris to “inflame” the local microenvironment, altering the NK cell cytotoxic modes, i.e., necroptosis vs. apoptosis, by modulating target membrane dynamics can enhance or damp the secondary inflammatory response subsequent to NK cell killing. This may hold the potential of, e.g., amplifying the anti-tumor effect of adoptive cell transfer therapy beyond primary NK cell killing.

Our result that membrane dynamics determine sensitivity to necroptotic killing has interesting implication for cancer, and perhaps for the role of necroptosis in homeostasis and disease more broadly. The Rho family of GTPases, e.g., Rac1 and the associated kinases, e.g., ROCK, are involved in key oncogenic signaling pathways, particularly by activating the actomyosin network and promoting cancer invasion and metastasis [33, 34]. Up-regulation of Rac1 activity has been frequently observed in human cancer [35] and Rac1 inhibitor, such as NSC23766 that we used in this study, is being actively pursued as anticancer therapeutic. Our data showed that tuning membrane dynamics by targeting Rac1 and ROCK is an effective strategy to influence the cytotoxic mode of NK cells. For cancers with elevated activity of Rac1 and/or Rho/ROCK signaling, they are likely more sensitive to proinflammatory killing by NK cells via necroptosis. Alternatively, inhibitors of the RTK signaling pathways that block protrusive and contractile membrane activity could be possibly employed to promote apoptotic killing in NK cell therapy. Necroptosis has been implicated as a major tissue damage pathway in diverse diseases, including stroke and neurodegeneration [36, 37]. Previously characterized necroptosis is typically activated by death receptor in combination with caspase inhibition [31]. Our finding that necroptosis can be triggered by granzyme and alteration in the actomyosin network and cell cortex may also be involved provide a novel mechanistic angle to investigate alternative regulatory pathway(s) underlying necroptosis in other disease contexts.

In terms of the mechanism underlying NK cell-induced necroptosis, our results suggested that a highly dynamic membrane, e.g., in MCF7 cells, likely facilitates excessive lytic granule transfer and subsequently results in rapid induction of high-level granzyme activities. Such high level of activated granzymes may preferentially cleave substrates associated with the actomyosin network, e.g., the ERM family proteins (actin-membrane linkers) [38], leading to actin depolymerization and collapse of the cell cortex. However, it is unclear whether the necroptosis machinery is activated due to the disruption of actomyosin network and cell cortex or is activated directly by the proteolytic substrates of granzymes. Yet to be resolved is also identity of the specific granzyme responsible for the NK cell-induced necroptosis. The majority of necrotic MCF7 cells that we observed did not display a significant loss of the GrzmB-FRET signal before the onset of necrotic blebs, which may suggest that the rapid necroptotic cytotoxicity is triggered not by granzyme B, but an alternative granzyme in NK cells, e.g., granzyme A or granzyme M.

Previous studies have showed Ca^2+^ played an important role in mediating lytic granule transfer upon NK-target cell interaction. We confirmed that chelating Ca^2+^ by EGTA abrogated both granzyme-B-dependent apoptosis and necrosis induced by primary NK cells. However, we did not observe high Ca^2+^ concentration shifted NK cell killing to the necrosis mode, as reported by Backes et al [15]. Increasing Ca^2+^ concentration up to 3.5 mM did not enhance necrotic killing of SMMC-7721 or U-2 OS cells. And Ca^2+^ concentration higher than 3.5 mM triggered cell death in the absence of NK cells, possibly due to the high salt concentration and osmotic stress. Given that Backes et al used suspension cell lines as NK cell targets, e.g., Jurkat E6-1 and K562, while we used epithelial targets, the difference between our results regarding the role of Ca^2+^ in necrotic killing could be due to target cell type variation, i.e., suspension vs. adherence, which is of interest for further study.

## Materials and Methods

### Isolation of primary NK cells from human blood

Primary human NK cells were isolated from fresh human buffy coat (within 24 hours of blood donation) obtained from the Hong Kong Red Cross. The buffy coat was first separated by Ficoll-Paque Plus solution and the NK cells were then purified from the Peripheral Blood Mononuclear Cells (PBMCs) by negative selection using the EasySep Human NK Cell Enrichment Kit (STEMCELL Technologies), according to manufacturer’s protocol. Only NK cell prep with purity higher than 90% (measured by FACS analysis of CD56 staining) was used for experiments. Fresh human NK cells were cultured for 3 days at a density of 1×10^6^ cells/ml in RPMI 1640 medium (Gibco, Thermo Fisher) containing 10 ng/ml recombinant human IL-2 (Gibco, Thermo Fisher), 10% heat-inactivated Fetal Calf Serum (Gibco, Thermo Fisher), 100 U/ml penicillin and 100 μg/ml streptomycin (Gibco, Thermo Fisher), prior to experiments.

### Cell lines and cell culture

The live-cell Förster Resonance Energy Transfer (FRET) construct that reports on the granzyme-B proteolytic activity as well as the mitochondrial reporter, IMS-RP, were engineered into the five selected epithelial cell lines by retroviral infection, as described previously [20, 21]. Isogenic clones of the fluorescent reporter cell lines that exhibited cell death response to primary NK cell killing most similar to their respective parental lines were selected for performing the live-cell imaging assays. Cell lines were cultured in appropriate medium supplemented with 10% Fetal Calf Serum, 100 U/ml penicillin and 100 μg/ml streptomycin. Specifically, U-2 OS was maintained in McCoy’s 5A (Modified) Medium; MCF7 was maintained in RPMI 1640 Medium; SMMC-7721, HeLa and LO2 were maintained in Dulbecco’s modified Eagles Medium (DMEM). For all co-culture experiments with primary NK cells, the NK-to-target cell ratio of 3:1 were used and 50 ng/ml IL-2 was supplemented in the medium. To image the dynamics of F-actin, we transiently transfected MCF7 and SMMC-7721 cells with the Utrophin-GFP reporter (Plasmid #26737, Addgene) using X-tremeGENE HP DNA Ttransfection Reagent (Roche). Time-lapse imaging was performed 48-72 hour after the transfection.

### Chemicals and neutralizing antibodies

The following inhibitors were purchased from Selleck Chem: NSC23766 (Rac1 inhibitor; used at 120 μM for MCF7 and 50 μM for SMMC-7721), necrostatin-1 (RIP1 inhibitor; used at 100 μM), GSK’872 (RIP3 inhibitor; used at 5 μM) and necrosulfonamide (MLKL inhibitor; used at 10 μM). Y27632 (ROCK inhibitor; used at 5 μM), Concanamycin A (Vacuolar type ATPase inhibitor; used at 10 nM) and 3,4-Dichloroisocoumarin (granzyme inhibitor; used at 40 μM) were purchased from Sigma, Santa Cruz Biotechnology and Cayman Chemical, respectively. Cells were pre-treated with the inhibitors for 2 hours before experiments, except for NSC23766 (5-12 hour pre-treatment), Concanamycin A (4-hour pre-treatment) and 3,4-Dichloroisocoumarin (20-minute treatment and then the inhibitor was removed to prevent toxicity to NK cells in the co-culture experiments). To block the activity of various NK cell receptors, the following neutralizing antibodies from Biolegend were used: Purified antihuman HLA-A,B,C antibody (Clone W6/32), CD94 (Clone DX22,), NKG2D (Clone 1D11), NKp46 (Clone 9E2), and DNAM-1 (Clone 11A8). Except for HLA-A,B,C antibody, which was used at 5 μg/ml, the rest of the antibodies were used at 10 μg/ml.

### Gene knockdown by RNAi

siRNA oligos for knocking down RIP1 (5’-CCACUAGUCUGACGGAUAA-3’), RIP3 (5’-CCAGAGACCUCAACUUUCA-3’) and MLKL (5’-CAAACUUCCUGGUAACUCA-3’) were custom synthesized by Dharmacon. Dharmacon On-Target plus siControl (#D-001810-01) was used as non-targeting siRNA control. RIP1 siRNA oligo was used at 40 nM, RIP3 siRNA at 60 nM, and MLKL at 30 nM. siRNA transfections were performed using Lipofectamine (Thermo Fisher) according to manufacturers’ instructions. Experiments were conducted after 48 hours of gene silencing. Gene knockdown efficiency was probed by western blot using the following primary antibodies from Cell signaling: RIP1 (#3493), RIP3 (#13526), and MLKL (#14993).

### Time-lapse microscopy and image analysis

Cells were plated in 96-well μ-plate (ibidi), and cultured in CO_2_-independent medium (Gibco, Thermo Fisher) supplemented with 10% heat-inactivated FCS, 4 mM L-glutamine, 100 U/ml penicillin and 100 μg/ml streptomycin. Cells were imaged either by a Nikon Eclipse TE-2000 inverted microscope enclosed in a humidified chamber maintained at 37°C with a 20X plan Apo objective (NA = 0.95), or by the Andor Dragonfly confocal microscope fitted with an incubation chamber. Images were acquired every 30 seconds to 4 minutes and data were viewed and analyzed using the NIS software (Nikon) or Imaris. Target cell death was scored morphologically by blebbing followed by cell lysis, and the time from NK cell addition to morphological target cell death was plotted as cumulative survival curves. The onset of apoptosis is scored by an abrupt transition from punctate to smooth localization of the mitochondria reporter, IMS-RP. To quantify the time course of the GrzmB-FRET signal, ImageJ was used to segment the individual cells and measure the total intensity of YFP and CFP signal, respectively.

### Flow cytometry analysis of surface ligand and receptor expression

1×10^6^ epithelial target cells were washed and re-suspended in the cell staining buffer (Biolegend) containing different dye-conjugated primary antibodies at 1 μg/ml or 5 μg/ml for 30 minutes at 4°C. Cells were then washed and stained with the Zombie NIR (cell death marker, Biolegend). The stained cells were analyzed by flow cytometry with a FACSCanto II cytometer (BD Biosciences). Primary antibodies targeting the various surface ligands and receptors as well as the corresponding control include: FITC anti-human HLA-A,B,C (Clone W6/32, Biolegend), FITC anti-human ICAM-1/CD54 (Clone DX2, Biolegend), FITC anti-human Fas/CD95 (Clone DX2, Biolegend), APC anti-human Nectin-2/CD112 (Clone TX31, Biolegend), APC anti-human PVR/CD155 (Clone SKII.4, Biolegend), PE anti-human MICA (Clone 159227, R&D Systems), PE anti-human MICB (Clone 236511, R&D Systems), PE anti-human ULBP-2/5/6 (Clone 165903, R&D Systems), FITC Mouse IgG1 κ Isotype Control (Clone MOPC-21, Biolegend), APC Mouse IgG1 κ Isotype Control (Clone MOPC-21, Biolegend), and PE Mouse IgG2b κ Isotype Control (Clone MPC-11, Biolegend).

## Supplementary Materials

Supplementary materials of this article include:

**Movie S1.** Time-lapse movie of MCF7 cell expressing the GrzmB-FRET reporter (green and blue) in co-culture with primary NK cells stained by lysobrite (red). The image is the overlay of the GzmB FRET reporter, lysobrite and bright-field signals. Time is indicated in the unit of hour:minute.

**Movie S2.** Time-lapse movie of SMMC-7721. The image is the overlay of the GzmB FRET reporter, lysobrite and bright-field signals.

**Movie S3**. Confocal live-cell imaging of F-actin (green) in MCF7 cell in co-culture with NK cells stained with lysobrite (red).

**Movie S4**. Confocal live-cell imaging of F-actin (green) in SMMC-7721 cell in co-culture with NK cells stained with lysobrite (red).

## Acknowledgements

We thank Dr. Paul Choi and Dr. Timothy Mitchison (Department of Systems Biology, Harvard Medical School) for the granzyme-B FRET reporter construct, and Dr. John Albeck and Dr. Peter Sorger (Department of Systems Biology, Harvard Medical School) for the IMS-RP retroviral vector. This work was supported by the Hong Kong Research Grant Council (#12101514, #T12-710/16-R and #C2006-17E) to J. Shi. The authors declare no conflict of interest.

## Author contributions

J.S. designed and supervised the study; Y.Z. performed the experiments; Y.Z., J.X. and J.S. analyzed the data; J.S. and Y.Z. wrote the paper.

## Notes

### Competing Interest Statement

The authors have declared no competing interest.

